# Tocilizumab induces significant changes in longitudinal proteomes of blood serum from patients with severe COVID-19 pneumonia

**DOI:** 10.64898/2026.05.05.723025

**Authors:** José Cordero, Graciela Bravo, Pedro H. Silva, Benjamin Lozano, Elizabeth Rivas, Valeria Labra, Daniela Villalobos, Pablo Saldivia, Mauricio Hernandez, Elard S. Koch, Cristian Vargas, Estefania Nova-Lamperti, Nelson P. Barrera, Jaime Retamal

**Affiliations:** Faculty of Biological Sciences, Pontificia Universidad Católica de Chile, Santiago 8331150, Chile; Department of Intensive Medicine, Faculty of Medicine, Pontificia Universidad Católica de Chile, Santiago 8330077, Chile; Fundación de Investigación San Ramón (FISAR), San Pedro de la Paz, Concepcion 4030563, Chile; Programa de Doctorado en Biotecnología Molecular, Facultad de Ciencias Biológicas, Universidad de Concepción, Concepcion 4070386, Chile; MELISA Institute, Concepción 4130000, Chile; Molecular and Translational Immunology Laboratory, Department of Clinical Biochemistry and Immunology, Faculty of Pharmacy, Universidad de Concepción, Concepcion 4070409, Chile

## Abstract

Coronavirus disease 2019 (COVID-19) shows highly variable clinical outcomes that are not fully explained by age or comorbidities, underscoring the importance of host molecular responses in determining disease severity. Proteomic and multi-omics studies have linked severe COVID-19 to profound dysregulation of immune, inflammatory, and coagulation pathways, and have shown that circulating protein signatures can predict clinical trajectories. Tocilizumab (TCZ), a monoclonal antibody targeting the interleukin-6 receptor (IL-6R), is an established therapy for IL-6–driven inflammatory diseases and can normalize aberrant molecular profiles. Here, we applied longitudinal serum proteomics to patients with severe SARS-CoV-2 pneumonia treated with TCZ to further characterize how IL-6R blockade reshapes the systemic inflammatory milieu. After TCZ administration, several clinical and inflammatory markers, including C-reactive protein (CRP), CCL5 and CXCL10, decreased. Proteomic profiling revealed that TCZ exerts a sustained effect on the serum proteome, with the most pronounced changes emerging 7 days after treatment. These changes were associated with a broad reconfiguration of the proteomic profile toward a pattern resembling a healthy physiological state, characterized by the restoration of key protein abundances to levels comparable to those observed under homeostatic conditions. Collectively, our findings support that TCZ treatment contributes to the normalization of the inflammatory state in severe COVID-19 and represents a viable therapeutic option for managing the acute inflammatory phase of the disease, while also highlighting additional pathways and biomarkers involved in this recovery process.

## Introduction

Coronavirus disease 2019 (COVID-19) caused by severe acute respiratory syndrome coronavirus 2 (SARS-CoV-2) had spread globally causing several millions of deaths worldwide. This pandemic stressed healthcare systems and imposed substantial social and economic costs on a global scale, making it one of the most significant public health crises in modern history (Huang et al., 2020; The Lancet Microbe, 2021). One of the main characteristics of COVID-19 is its highly variable clinical presentation, which ranges from asymptomatic infection to critical illness characterized by acute respiratory distress syndrome (ARDS), multiorgan dysfunction, and death (Di et al., 2020; The Lancet Microbe, 2021). This variability is only partially explained by demographic or comorbidity-related risk factors such as age, obesity, and cardiovascular disease (Parotto et al., 2023), suggesting that host molecular responses play a decisive role in the disease progression.

Large-scale proteomics and multi-omics studies have deepened into these factors by characterizing the molecular trajectories of COVID-19 patients over the progression of the disease, identifying dysregulations of immune, inflammatory, and coagulation pathways in patients with severe disease, such as elevated levels of C-reactive protein (CRP), interleukin-6 (IL-6), D-dimer, ferritin, and markers of neutrophil degranulation (Babačić et al., 2023; Demichev et al., 2021; Filbin et al., 2021; Messner et al., 2020). Also, high-throughput proteomic platforms have revealed protein signatures reflecting complement activation, endothelial injury, acute-phase response, and metabolic reprogramming that differentiate mild from severe cases (Babačić et al., 2023; Demichev et al., 2021; Filbin et al., 2021; Messner et al., 2020). Longitudinal plasma proteome profiling has demonstrated that these signatures not only correlate with clinical severity but also predict patient trajectories and therapeutic needs with high accuracy (Babačić et al., 2023; Demichev et al., 2021; McArdle et al., 2021).

Progression from mild or moderate COVID-19 to severe or critical illness is largely attributed to an imbalanced or dysregulated host immune response, rather than to uncontrolled viral replication. Upon SARS-CoV-2 infection, the innate immune system initiates a rapid antiviral defense via interferons, chemokines, and pro-inflammatory cytokines, this initial response is essential for viral containment, but it can become pathogenic when it is excessive or prolonged, as in severe COVID-19. Here, this is established as a systemic inflammatory response syndrome (SIRS)-like condition, where myeloid cells are overactivated, hypersecretion of cytokines, and tissue-destructive immune pathways are present (Dritsoula et al., 2022; Gómez-Rial et al., 2020; Ogata et al., 2019; Tang et al., 2022).

Tocilizumab (TCZ) is a humanized monoclonal antibody that blocks both membrane-bound and soluble forms of the interleukin-6 receptor (IL-6R), in that way inhibiting IL-6–mediated pro-inflammatory signaling (Gordon et al., 2021). It was originally developed for autoimmune diseases, but now TCZ is approved for the treatment of rheumatoid arthritis (RA), juvenile idiopathic arthritis, giant cell arteritis and for cytokine release syndrome (CRS) associated with CAR T-cell therapies (Huang et al., 2020; Ogata et al., 2019). Its efficacy in these conditions is based on its ability to suppress IL-6–driven inflammation, reduce acute-phase reactants such as CRP, and modulate the immune response at both systemic and molecular levels (Yang et al., 2021). Tasaki et al. performed a longitudinal multi-omics analysis in RA patients treated with biological or targeted DMARDs (targeted synthetic therapies), including TCZ, and found that patients exhibited significant normalization of transcriptomic and proteomic profiles towards a healthy baseline (Tasaki et al., 2018). Such findings indicate that TCZ’s effects extend beyond symptom control and can be linked with the reprogramming of the systemic inflammatory landscape. These insights support the notion that plasma proteome remodeling could occur in COVID-19 patients following TCZ treatment.

Therefore, we hypothesize that the use of TCZ in patients with severe SARS-CoV-2 pneumonia is associated with an early modification of their molecular serum phenotype from a “severe” to a “mild” profile. To confirm this hypothesis, we determined the serum proteome of patients with severe SARS-CoV-2 pneumonia before and after administration of TCZ, and compared the findings with previously published molecular markers in patients with different degrees of COVID-19 severity. The results showed that after TCZ treatment, some clinical and inflammatory markers such as CRP, CCL5 and CXCL10 decreased. Serum proteomics showed that TCZ induces a long-term effect in the identified proteins, where the main changes were detected 7 days after treatment. These changes led to a reconfiguration of the proteomic profile toward a pattern resembling a healthy physiological state, characterized by the restoration of key protein levels to values comparable to those observed under homeostatic conditions. Our findings support the conclusion that TCZ treatment contributes to the normalization of the inflammatory state in patients and represents a viable therapeutic option for managing the acute symptoms of COVID-19.

## Materials and Methods

### Study design and participants

This study explores longitudinal changes in the blood serum proteome of seven consecutive COVID-19 patients (IDs P1-P7) diagnosed with severe pneumonia and admitted to the Intensive Care Unit of UC-Christus Clinical Hospital on May 2020. Proteomic profiles were analyzed before and after administration of a single intravenous dose of TCZ (8 mg/kg). An additional healthy control group of seven individuals (IDs 1-7) was added (40 years old average, range 35-48).

### Data collection

Clinical and epidemiological data, along with blood serum samples, were collected at three time points: one day before the treatment (D0), and on days 1 (D1), 4 (D4) and 7 (D7) post-treatment. The study was approved by the Institutional Ethical Review Board of the Pontificia Universidad Católica de Chile (Protocol #201109002).

### Proteomic analysis

Blood serum samples from patients were subjected to high-abundance protein depletion using the Multiple Affinity Removal Spin Cartridge Human 14 (Agilent, USA), with 800 µg of native serum proteins loaded per column, following the manufacturer’s protocol. The flow-through containing the depleted protein fraction was precipitated using cold 100% acetone at a 5:1 (v/v) ratio and incubated overnight at −20 °C. Samples were then centrifuged at 15,000 × g for 10 minutes, the supernatant was discarded, and the resulting pellet was washed three times with 90% (v/v) acetone. The protein pellets were dried using a rotary vacuum concentrator at 4 °C and subsequently resuspended in 8 M urea with 25 mM ammonium bicarbonate (pH 8.0).

Proteins were reduced with 20 mM dithiothreitol (DTT) for 1 hour at room temperature, followed by alkylation with 20 mM iodoacetamide for 1 hour in the dark. Protein concentration was determined using Qubit Protein Quantification Kit (Thermo Fisher Scientific), and 10 µg of total protein was used for digestion. Samples were diluted to a final concentration of 1 M urea with 25 mM ammonium bicarbonate (pH 8.0) and digested overnight at 37 °C using sequencing-grade trypsin/Lys-C (Promega) at a 1:50 enzyme-to-substrate ratio. Peptides were purified using Sep-Pak Vac C18 cartridges (Waters, USA), according to the manufacturer’s instructions. Eluted peptides were dried using a rotary vacuum concentrator at 4 °C and resuspended in 2% acetonitrile (ACN) with 0.1% formic acid (FA) (Merck, Germany). Final peptide concentrations were measured using the Direct Detect spectrometer (Merck Millipore).

High pH reversed-phase fractionation was performed on an AKTA Avant 25 (General Electric) coupled to a refrigerated fraction collection. Purified peptides were separated on a reversed-phase column BHE 2.1 cm x 5 cm (Waters) at a flow rate of 0.2 mL/min at pH 10. The binary gradient started from 3% buffer B (90% ACN in 5 mM Ammonium Formate pH 10), followed by linear increases to first 40% B within 30 min, to 60% B within 15 min, and finally to 85% B within 5 min. Each sample was fractionated into 24 fractions in 400 µl volume intervals. The fractions were dried in a vacuum-centrifuge and reconstituted in water with 2% ACN and 0.1% FA and concatenated in 8 fractions.

### Liquid chromatography

All peptide fractions were analyzed using a nano-ultrahigh-performance liquid chromatography system (nanoElute, Bruker Daltonics) coupled to a nano-flow setup. Peptides (200 ng per injection) were separated on an Aurora Series C18-fused silica column (75 μm inner diameter, 360 μm outer diameter, 25 cm length, 1.6 μm particle size) with an integrated emitter tip (IonOpticks, Australia). Separation was performed at 50 °C using a 90-minute gradient at a flow rate of 400 nL/min. Mobile phase A consisted of water with 0.1% formic acid (FA), and mobile phase B was acetonitrile with 0.1% FA. The gradient was programmed as follows: 2% to 17% B over 37 minutes, 17% to 25% B over 15 minutes, and 25% to 35% B over 8 minutes, followed by a high-organic wash at 85% B and re-equilibration.

### The timsTOF Pro Mass Spectrometer

All peptide fractions were analyzed using a hybrid trapped ion mobility spectrometry (TIMS) quadrupole time-of-flight (QTOF) mass spectrometer (timsTOF Pro, Bruker Daltonics), equipped with a CaptiveSpray nano-electrospray ion source. The mass spectrometer was operated in data-dependent acquisition (DDA) mode to enable ion mobility-enhanced spectral library generation. Both the accumulation and ramp times were set to 100 ms, and mass spectra were acquired in positive electrospray mode over an m/z range of 100–1700. Ion mobility separation was performed across a range of 0.6 to 1.6 Vs/cm². Each acquisition cycle lasted 1.16 seconds and included one full TIMS-MS scan followed by 10 parallel accumulation–serial fragmentation (PASEF) MS/MS scans.

### Database searching

Tandem mass spectra were extracted by Tims Control version 2.0. Charge state deconvolution and deisotoping were not performed. All MS/MS spectra were analyzed with search engine PEAKS Studio X+ (Bioinformatics Solutions Inc, Canada version 10.5 (2019-11-20)) against the SwissProt human proteome database, assuming trypsin as the digestion enzyme. The search parameters were set to a parent ion mass tolerance of 50 ppm and a fragment ion mass tolerance of 0.050 Da. Carbamidomethylation of cysteine was specified as a fixed modification, while deamidation of asparagine and glutamine, oxidation of methionine, N-terminal acetylation, and carbamylation of lysine and the protein N-terminus were included as variable modifications.

### Statistical analysis

Data analysis and statistical testing were performed using the R environment (version 4.3.1;R Core Team, www.R-project.org). Missing values were imputed with zero/minimal value of the respective time-point. Raw intensities were transformed to log_2_ and then scaled before Principal Component Analysis (PCA). For differential expression analysis, fold-change (FC) was calculated for each value based on the mean/median of each time-point group. Differences in log₂-transformed intensities between time-point groups were assessed using a repeated measures eBayes test (from *limma* package in R).

For comparisons of biomarker expression between groups using boxplots, statistical differences were evaluated using the non-parametric Wilcoxon rank-sum test (*wilcox_test*, *rstatix* package). Proteins were considered differentially expressed if they had a p-value < 0.05 (Pascovici et al., 2016) and an absolute fold change (|FC|) greater than 1.

### Pathway Analysis

To identify enriched canonical pathways, biological functions, and disease associations from the proteomics data, the software QIAGEN IPA (QIAGEN Inc., https://digitalinsights.qiagen.com/IPA) was used. An Expression Core Analysis was performed on the uploaded dataset using a statistical threshold of p < 0.05 and an absolute log₂ fold change greater than 0.5 or smaller than -0.5. In order to complement these results, an expression analysis was carried out using Reactome and the camera algorithm based on the normalized proteomics data.

To assess temporal changes in protein expression and the predicted activation states of canonical pathways, a Comparison Analysis was carried out for the following time-point contrasts: D1 vs D0, D7 vs D0, and D7 vs D1.

## Results

In this study, patients diagnosed with COVID-19 and presenting severe pneumonia were recruited and treated with TCZ aiming to decrease disease severity. To assess the therapeutic impact of TCZ, clinical parameters were measured before and after drug administration to evaluate temporal changes. The results are shown in Figure 1. Recruited patients were 57 years old average (range 45-70) and TCZ was applied 8.5 (7.3-13 range) days after starting symptoms and within 1 day after ICU admission. While CRP decreased after 1 day of treatment with TCZ (Figure 1A), PAFI, ferritin, D-dimer and RAL did not change during the measured times (Figure 1 B-E).

**Figure 1.**
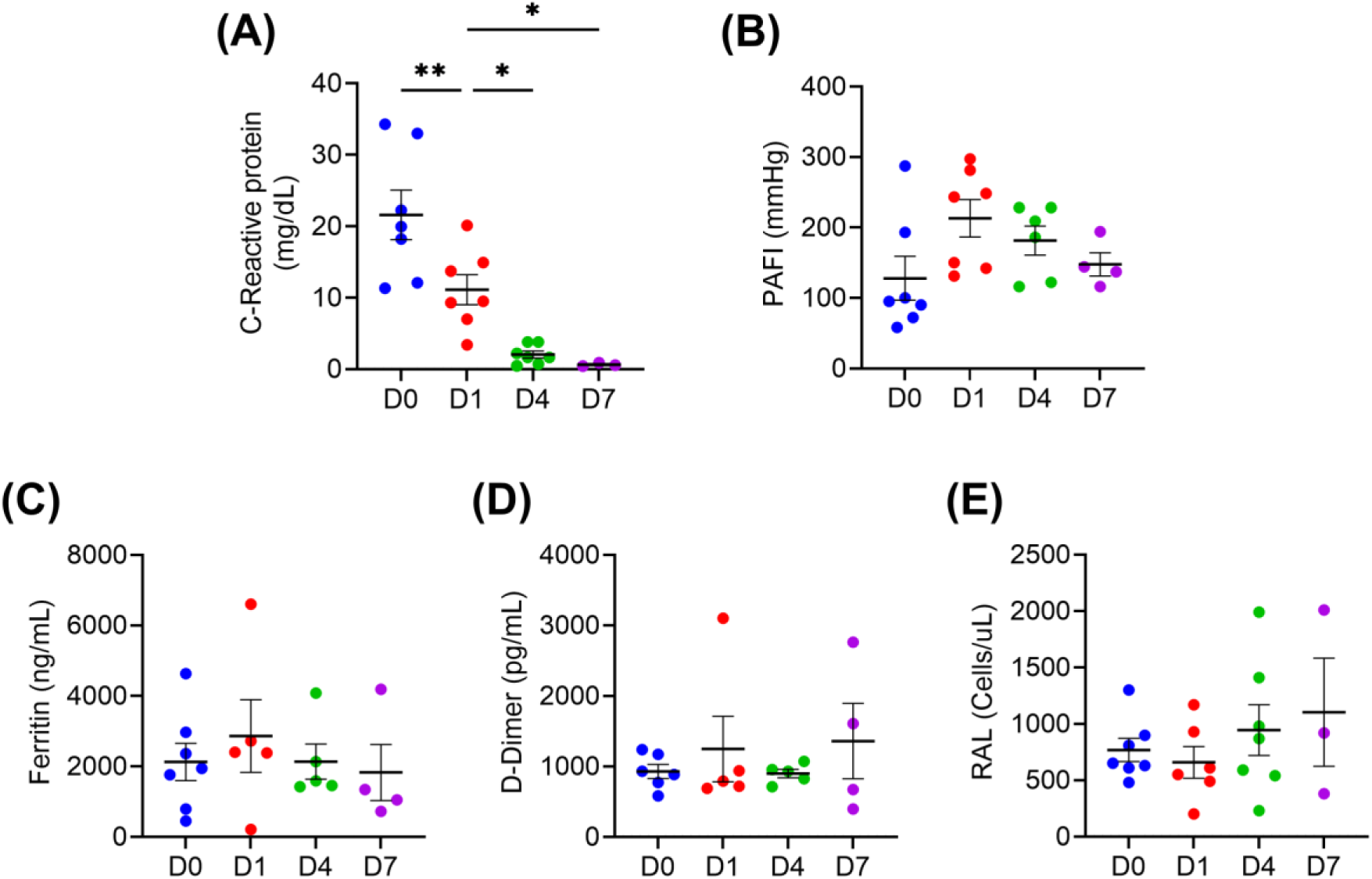
Clinical parameters of severe COVID-19 pneumonia patients. Clinical parameters were measured before and after drug administration to evaluate the changes. (A) C-reactive protein, (B) PaO2/FIO2 ratio, (C) ferritin, (D) D-dimer, and (E) RAL were measured in serum. One one-way ANOVA followed by Tukey’s multiple comparison test was performed to compare the data. n = 7; (*\*p<0.05, **p<0.01, ***p<0.001, ****p<0.0001*).

To evaluate changes in the inflammatory profile, serum cytokine levels were measured using multiplex assays at multiple time points and compared with values from the healthy control group (Figure 2). CCL5 levels remained elevated in COVID-19 patients compared to healthy controls despite TCZ treatment (Fig. 2A). Similarly, CXCL10 levels decreased by day 7 following TCZ administration but remained higher than in healthy individuals (Fig. 2B). No significant differences were observed in CXCL9 and CCL2 levels between patients and healthy controls (Fig. 2C,D). IL-8 levels showed a decreasing trend after TCZ treatment; however, they remained elevated relative to controls. (Figure 2E). Finally, IL-6 levels did not change following TCZ treatment and were comparable to those of healthy controls. These findings suggest that TCZ exerts anti-inflammatory effects, as evidenced by the post-treatment reductions in CRP, CCL5, and CXCL10 levels.

**Figure 2.**
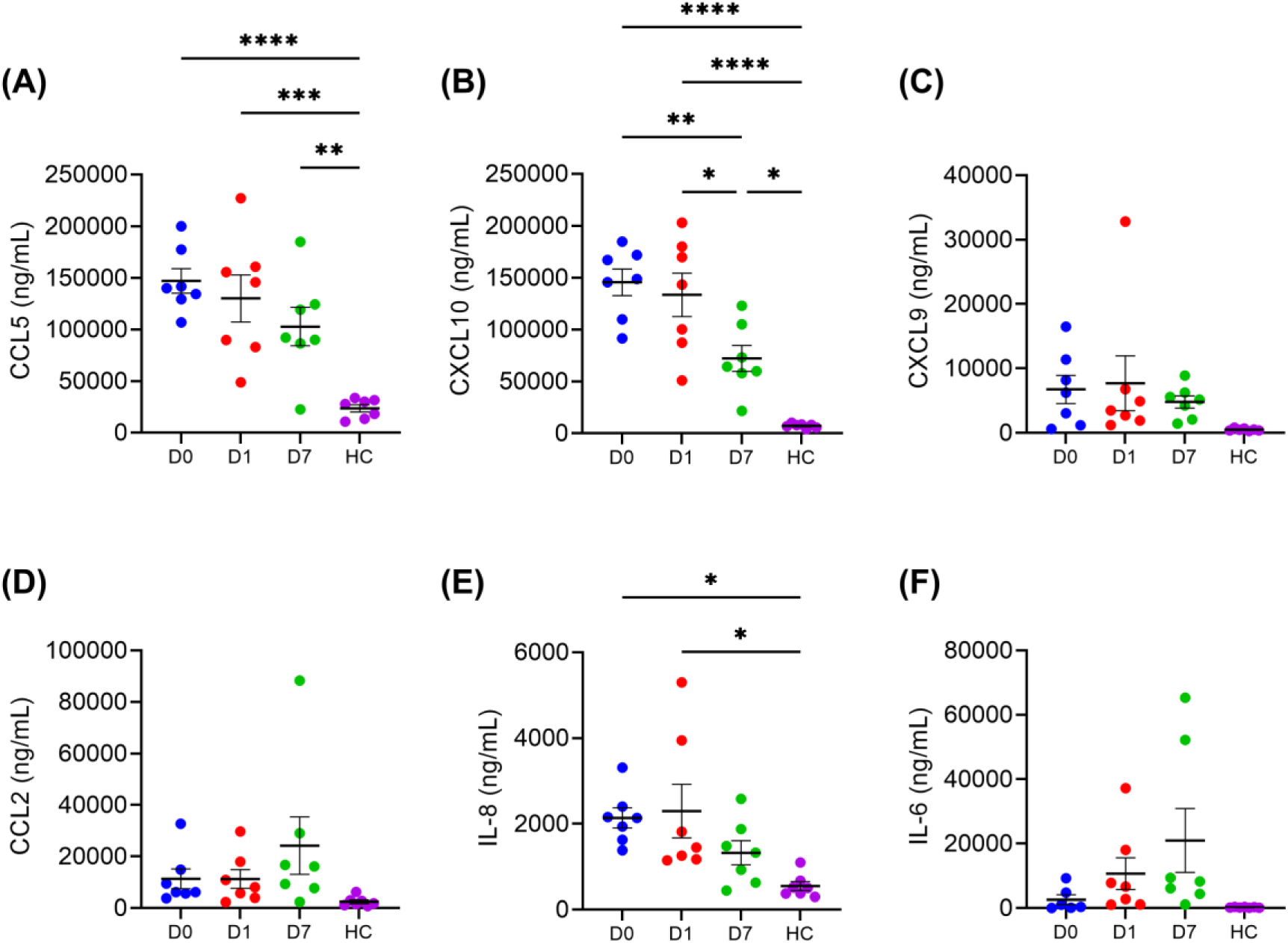
Cytokines and chemokines levels in serum of severe COVID-19 pneumonia patients and healthy control group. Cytokines and chemokines were measured before and after TCZ administration to evaluate the changes until day 7 in severe COVID-19 pneumonia patients. In addition, cytokines and chemokines were also measured in a healthy control group without TCZ (A) CCL5, (B) CXCL10, (C) CXCL9, (D) CCL2, (E) IL-8, and (F) IL-6 levels were measured by multiplex. One one-way ANOVA followed by Tukey’s multiple comparison test was performed to compare the data. n = 7; (*\*p<0.05, **p<0.01, ***p<0.001, ****p<0.0001*).

To assess how TCZ treatment alters the plasma proteome of COVID-19 patients, a comprehensive deep proteomic analysis was conducted. To evaluate if changes in blood proteome were associated with a reduced severity of the disease, the severity of biomarkers was evaluated as previously described by Shen et al., Messner et al. and Overmyer et al. (Messner et al., 2020; Overmyer et al., 2020; Shen et al., 2020) (Figure 3A, B). The literature review yielded 204 candidate proteins, of which 119 are enriched in severe disease (“UP” markers) and 85 are reduced (“DOWN” markers). Mapping these candidates onto our spectral library, 179 biomarkers in total—99 UP and 80 DOWN, were identified and quantified. Overall, in the category ‘UP’ a reduction is shown in the protein levels after TCZ treatment (comparisons D1 vsD0 and D7 vsD0) particularly in the grouped proteins 2,while an increase is shown in the levels of the proteins in the category ‘DOWN’ at D7 vs D0 after TCZ treatment particularly in the grouped proteins 1 and 3, indicating that IL-6 receptor blockade shifts the circulating proteome toward a profile associated with milder disease.

**Figure 3.**
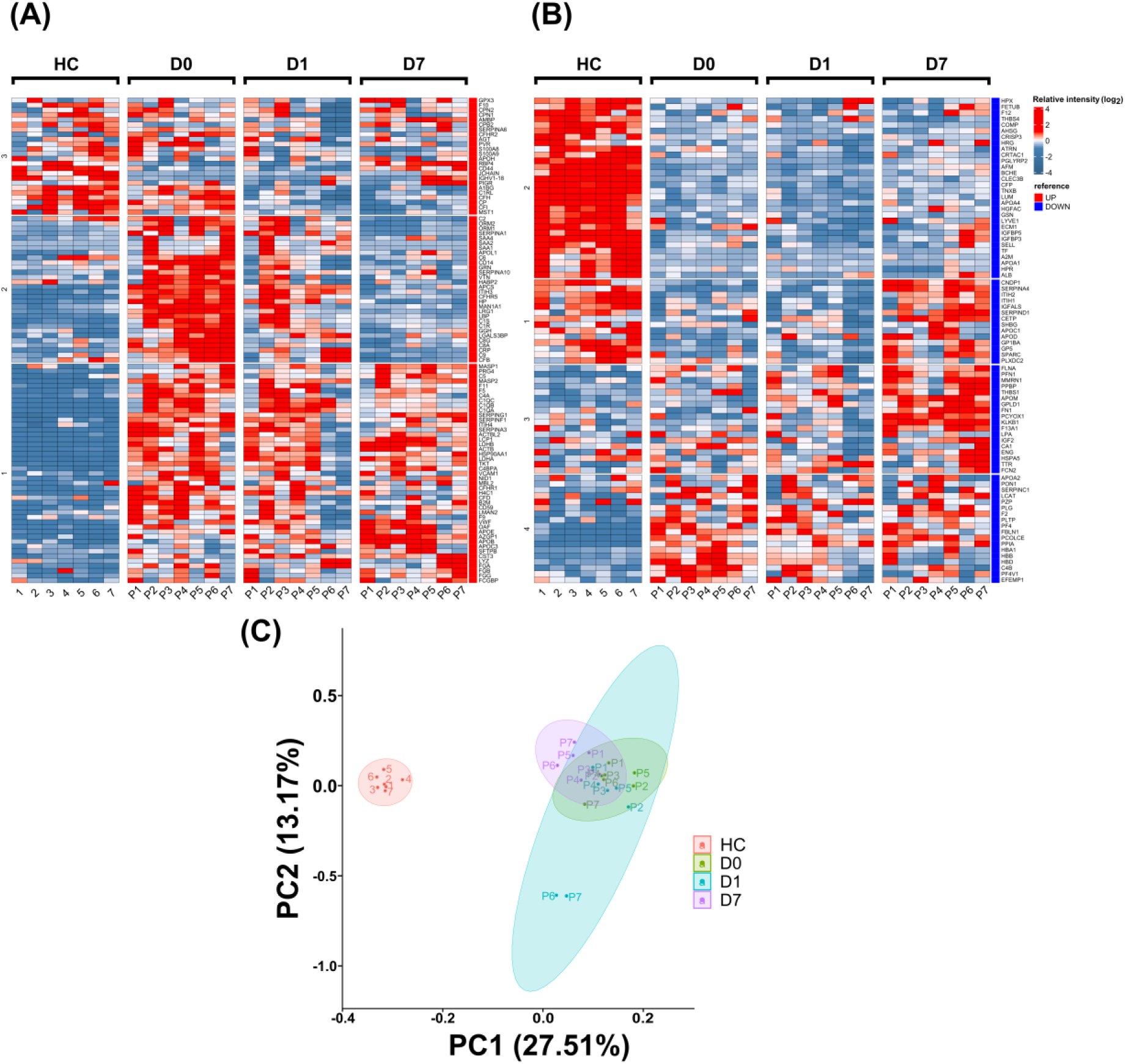
Longitudinal blood-serum proteomes of severe COVID-19 pneumonia patients after TCZ treatment. Heat map representations for a number of known up-regulated (A) and down-regulated (B) protein biomarkers previously determined in severe COVID-19 patients (Messner et al., 2020; Overmyer et al., 2020; Shen et al., 2020) present in 7 COVID-19 patients with severe pneumonia treated with TCZ. In addition, protein groups were analyzed via hierarchical clustering as indicated by numbers (1 to 4) at the left side. Each heat map begins with the profile of healthy controls (HC, n = 7) as a baseline reference, followed by samples from the seven severe COVID-19 patients on days 0, 1, and 7 (D0, D1, D7) after TCZ therapy. Relative protein signal intensity is shown as log₂ values; higher intensities appear in red and lower intensities in blue. Biomarkers are ordered according to their known tendency to rise or fall with worsening disease. (C) Principal component analysis (PCA) of differentially expressed proteins (DEPs) demonstrates clear separation between healthy controls and patient samples, as well as the trajectory of each patient from D0 through D7, highlighting treatment-induced shifts toward the healthy proteomic profile.

Deep proteomics analysis identified an average of 1,564 serum proteins per sample (range 1,112–2,110), from which 552 differentially expressed proteins (DEPs) were selected for downstream analysis. DEPs were identified based on pairwise comparisons between time points: D1 vs. D0, D7 vs. D0, and D7 vs. D1. Principal component analysis (PCA) revealed a temporal shift in the patients’ serum proteomic profiles from baseline (D0) to Day 7 (D7), with less overlapping between D0 and D7 (Figure 3C). Volcano plots illustrating the differential expression patterns for each of these comparisons are provided in Supplementary Figure 1. Interestingly, once graphed the PCA for the healthy control it is observed a shorter centroid-to-centroid distance with the D7 data (28.75 for D7 vs HC, 31.91 for D1 vs HC and 32.47 for D0 vs HC).

Building upon these global proteomic shifts, Figure 4A presents a heatmap depicting the expression log ratios of the top 25 most DEPs across three time-point comparisons following TCZ treatment in patients with severe COVID-19. A pronounced downregulation is observed in early acute-phase response proteins—SAA1, SAA2, CRP, ORM1, ORM2, and SERPINA1—which show large negative log ratios, particularly between D7 vs D0 and D7 vs D1. For instance, SAA1 shows a dramatic decrease of −5.27 at D7 vs D0 and −4.78 at D7 vs D1, consistent with the known suppression of IL-6–mediated inflammatory pathways targeted by TCZ (Beyer et al., 2011; Loricera et al., 2015). Similarly, SAA2 is downregulated with values of −4.02 and −3.29, and CRP drops to −3.33 and −2.61 at the same comparisons.

**Figure 4.**
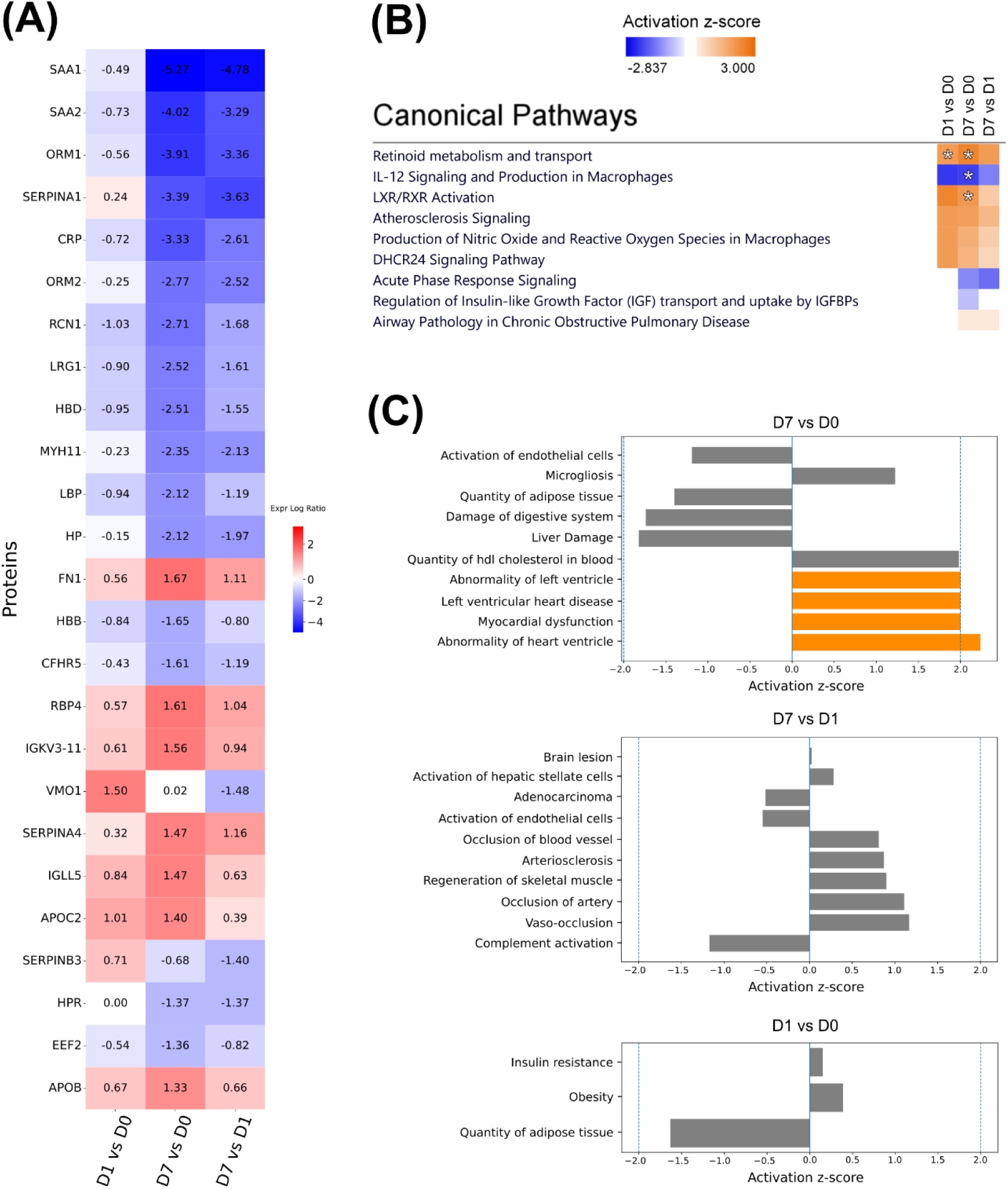
COVID-19 proteomic heat map of top 25 differentially expressed proteins. (A) Heat-map showing the top 25 proteins that are significantly (p < 0.05) and differentially expressed. Biomarkers are classified as up-regulated when the log₂ fold-change (log₂FC) > 0.5 and down-regulated when log₂FC < 0.5. The numbers within the cells correspond to the sum of the absolute log₂FC values for every observation of each biomarker. (B) IPA comparison analysis of canonical pathways across each time-point comparison. Bars are colored according to their z-score: orange indicates predicted activation, while blue indicates negative z-score, asterisks indicate |z-score| > 2 and p-value < 0.05. (C) IPA enriched biological functions and diseases for time-point comparisons D7 vs D0 and D7 vs D1, bars are colored according to their z-score: orange indicates a z-score > 2 while gray bars indicate |z-score| < 2.

Proteins such as RCN1, LRG1, HBD, HP, MYH11, and HBB also show consistent negative expression values (e.g., RCN1: −2.71, −1.68; HP: −2.12, −1.97) across time, indicating a broader anti-inflammatory effect post-TCZ treatment. In contrast, another cluster of proteins, notably IGKV3-11, IGLL5, SERPINA4, APOC2, RBP4, APOB, and FN1, display upregulation. FN1, for example, rises from 0.56 at D1 vs D0 to 1.67 at D7 vs D0, remaining elevated at 1.11 by D7 vs D1, suggesting a progressive increase over time. Likewise, IGKV3-11 and IGLL5 show robust upregulation peaking at D7 vs D0 with values of 1.56 and 1.47, respectively, reflecting possible compensatory or adaptive immune responses. These results are in concordance with the measure of standardized intensity (Supplementary Figure 2), where 7 days after TCZ treatment the levels of several severity markers are similar to the healthy controls.

In line with the temporal proteomic changes observed, Figure 4B displays a heatmap summarizing the predicted activation states, based on their z-score metric calculated by IPA, of key canonical pathways significantly modulated across the three comparisons: D1 vs D0, D7 vs D0, and D7 vs D1. Asterisks represent if the pathways meet the statistical threshold of a |z-score| > 2 and a p-value < 0.05 in order to be identified as significantly enriched and activated/inhibited.

Among the predicted inhibited pathways are Acute Phase Response Signaling and IL-12 Signaling and Production in Macrophages, particularly at early time points. These pathways remain suppressed through D7, indicating a persistent downregulation following TCZ treatment.

Conversely, pathways such as LXR/RXR Activation, Atherosclerosis Signaling, and Retinoid Metabolism and Transport show progressive activation, with increasing positive z-scores from D1 to D7. These pathways are involved in lipid metabolism, nuclear receptor signaling, and transport of small molecules, suggesting a metabolic shift occurring over the treatment period, this was also supported by the Reactome expression analysis that exhibited pathways related with lipid and cholesterol metabolism (Supplementary Figure 3). DHCR24 Signaling, associated with cholesterol biosynthesis and oxidative stress regulation, shows a strong positive z-score towards D7. At later stages, a negative z-score value of the Regulation of IGF Transport and Uptake by IGFBPs pathway is observed, suggesting a reduction in growth factor transport dynamics. Additionally, the Airway Pathology in Chronic Obstructive Pulmonary Disease pathway shows a positive trend at D7 vs D1.

Continuing the functional analysis, Figure 4C presents enriched biological functions and diseases from the Core Analysis performed by IPA on the protein expression profiles at D7 vs D0 (top panel), D7 vs D1 (middle panel) and D1 vs D0 (bottom panel). These bar plots display activation z-scores for significantly enriched functions and diseases, indicating either a predicted activation state “Increased” (orange) if the biological process has an activation z-score above 2, or “Decreased” (blue) in case of an activation z-score less than -2.

In the D7 vs D0 comparison, the highest activation z-scores were predominantly associated with cardiovascular-related disease and function annotations. IPA predicted an “Increased” state for multiple heart/ventricle terms, including abnormality of the heart ventricle (z-score = 2.236), myocardial dysfunction (z-score = 2.000), left ventricular heart disease (z-score = 2.000), and abnormality of the left ventricle (z-score = 2.000). In addition, quantity of HDL cholesterol in blood showed a strong positive trend toward activation (z-score = 1.982), while microgliosis also trended positively (z-score = 1.224), although it did not reach the threshold required for an “Increased” predicted state.

Conversely, IPA indicated inhibitory trends for liver damage (z-score = −1.822), damage of the digestive system (z-score = −1.739), quantity of adipose tissue (z-score = −1.400), and activation of endothelial cells (z-score = −1.191). Overall, these results suggest a dominant shift toward increased cardiovascular dysfunction and ventricular abnormality signatures at D7 relative to D0.

In the D7 vs D1 comparison, the highest positive activation z-scores were primarily associated with vascular-occlusive and cardiovascular-related annotations, led by vaso-occlusion (z-score = 1.167) and occlusion of artery (z-score = 1.109). Additional positive trends were observed for regeneration of skeletal muscle (z-score = 0.900), arteriosclerosis (z-score = 0.871), and occlusion of blood vessel (z-score = 0.810). More modest increases were noted for activation of hepatic stellate cells (z-score = 0.283), whereas brain lesion remained near baseline (z-score = 0.025).

Conversely, the strongest negative z-score corresponded to complement activation (z-score = −1.172), accompanied by inhibitory trends for activation of endothelial cells (z-score = −0.548) and adenocarcinoma (z-score = −0.514). Notably, none of the annotations reached the |z-score| ≥ 2 threshold, indicating that these changes reflect directional trends rather than high-confidence predicted increased or decreased states.

Finally, In the D1 vs D0 comparison, the pattern is dominated by a decreased trend for quantity of adipose tissue (z-score = −1.633). In contrast, obesity (z-score ≈ 0.391) and insulin resistance (z-score ≈ 0.152) show modest positive trends in activation z-scores.

Altogether, these functions indicate that by day 7, TCZ-treated patients exhibit a proteomic signature consistent with reduced inflammation-associated tissue damage and improved metabolic function, particularly in lipid and oxidative pathways, while some vascular remodeling signatures persist or begin to emerge in comparison to the earlier post-treatment time point.

## Discussion

COVID-19 has caused a profound global impact, with persistent effects manifesting as seasonal outbreaks and long-term sequelae in population (Parotto et al., 2023; The Lancet Microbe, 2021). In this study, we assessed whether TCZ administration could regulate the acute pneumonia caused by COVID-19 by evaluating both clinical parameters and longitudinal proteomic profiles in patient serum, focusing particularly on severity-associated biomarkers and signaling pathways.

To delineate the proteomic fingerprint most responsive to TCZ, we first generated a severity heatmap for a curated panel of severity biomarkers (Figure 3), which revealed a coherent trajectory aligned with clinical recovery, consistent with previous longitudinal proteomics studies (Messner et al., 2020; Overmyer et al., 2020; Shen et al., 2020).

At baseline, our cohort reproduced the canonical IL-6–dominant signature of severe COVID-19: key acute-phase reactants such as serum amyloid A (SAA1/2), C-reactive protein (CRP), haptoglobin (HP), α1-acid glycoproteins (ORM1/ORM2), and α1-antitrypsin (SERPINA1) were all markedly elevated (Demichev et al., 2021; Messner et al., 2020). Twenty-four hours post-TCZ administration, these proteins declined clearly and consistently, with further reductions by day 7, particularly SAA1/2, CRP, HP and ORM1/2, ultimately falling within or below the healthy reference range (Supplementary Figure 2). This pattern mirrors the “resolving” proteomic trajectory documented in survivors of critical illness (Filbin et al., 2021). Concurrently, HDL-associated and anti-inflammatory proteins rebounded: APOC1, APOC2, APOA4, and AZGP1 showed a sustained increase, while RBP4 and APOC4 increased in synchrony—indicative of HDL recovery previously linked to favorable outcomes (Babačić et al., 2023; McArdle et al., 2021). Our results also show that IL-6 does not present a significant change for patients before and after treatment with TCZ and healthy individuals (Figure 2), as reported previously (Aguareles et al., 2025).

This molecular profile was consistent with a predicted inhibition of the acute phase response pathway at D7, supported by both enrichment and predictive analyses using IPA (Figure 4), in line with previous inflammatory settings involving TCZ (Nakaok et al., 2013). In contrast, the LXR/RXR pathway was predicted to be activated at all-time points. Whereas prior studies indicate that COVID-19 suppresses this pathway (Tang et al., 2022), our findings align with TCZ’s recognized ability to restore anti-inflammatory signaling, as documented in conditions such as rheumatoid arthritis (RA) (Ogata et al., 2019). These results suggest that TCZ promotes sustained anti-inflammatory remodeling during the acute-to-resolution phase transition.

Beyond IL-6 modulation, TCZ influenced markers of endothelial function and complement activity. SERPINA4 (kallistatin) increased by >1 log₂, surpassing the healthy range. Elevated SERPINA4 has been independently associated with reduced microvascular damage and improved renal function (Demichev et al., 2022; Patel et al., 2024). Concurrently, pro-angiogenic LRG1 and the alternative complement amplifier CFHR5 declined, while fibronectin (FN1) increased by 1.7 log₂, collectively reflecting extracellular matrix remodeling and endothelial repair (Dritsoula et al., 2022). Soluble CD14 levels also declined, suggesting decreased monocyte activation (Gómez-Rial et al., 2020).

In parallel, IPA analysis revealed predicted activation of the nitric oxide (NO) and reactive oxygen species (ROS) production pathway in activated macrophages, despite TCZ’s overall anti-inflammatory influence. As NO and ROS are integral to host antiviral responses (Forman & Torres, 2002; MacMicking et al., 1997), their activation may reflect classical immune defense mechanisms against SARS-CoV-2 (Chernyak et al., 2020). However, caution is warranted, as TCZ has been associated with macrophage activation syndrome (MAS) in some pathological contexts (Tsuchida et al., 2017), suggesting that this activation may result from both host immunity and drug-induced immune dysregulation.

Cholesterol metabolism, another key axis in COVID-19 severity (Sohrabi et al., 2021), also emerged as relevant. The predicted activation of the DHCR24 pathway suggests compensatory lipid remodeling during viral clearance. Additionally, antiviral effects of cholesterol derivatives, such as inhibition of SARS-CoV-2 entry, have been documented (Murae et al., 2024), reinforcing the biological significance of these changes. These proteomic adjustments align with the upregulation of lipid-transport proteins observed in our study, supporting the notion of an anti-inflammatory metabolic recovery.

TCZ also mitigated cellular stress and hemolytic profiles: RCN1, an ER stress marker, dropped by −2.7 log₂, while free hemoglobin subunits HBD and HBB declined, consistent with restored membrane integrity and reduced hemolysis (Demichev et al., 2021; Filbin et al., 2021; Shen et al., 2020). Notably, predicted activation of the atherosclerosis signaling pathway—frequently exacerbated by viral infections (Chidambaram et al., 2024)—was observed consistently across timepoints. In RA cohorts, TCZ has shown no adverse effect on atherosclerotic progression (Moriyama et al., 2025), suggesting a vascular-protective dimension to its mechanism of action.

Further immunological recalibration was indicated by increased IGKV3-11, possibly reflecting neutralizing B-cell clonal expansion during convalescence (Wang et al., 2021). The activation of the retinoid metabolism pathway, implicated in immune homeostasis and IL-6 regulation (Nagpal et al., 1997; Rodan Sarohan, 2023), may represent both host response and TCZ-mediated modulation of the IL-6 axis. In contrast, the predicted inhibition of IL-12 signaling aligns with TCZ’s mechanism in attenuating cytokine storm responses (Gordon et al., 2021), particularly given that IL-6, IL-12, and TNF-α are major effectors in COVID-19–related hyperinflammation (Huang et al., 2020; Yang et al., 2021). Finally, periostin (POSTN) and SERPINB3 showed delayed or biphasic dynamics—early suppression followed by rebound—suggesting late fibroproliferative activity and possible organ-specific sequelae (Babačić et al., 2023).

Regarding the functional analysis, IPA predicted an increase in four cardiac-related functional categories: myocardial dysfunction, abnormality of the heart ventricle, left ventricular heart disease, and abnormality of the left ventricle. These functions and diseases were enriched and their activation states were driven by several proteins, with CKM (logFC = −0.616), CSF1 (logFC = −0.523), TNFRSF1B (logFC = −0.567), and CTSA (logFC = 0.517) contributing most strongly to the predicted increase, according to IPA.

CKM is an established cardiac biomarker involved in high-energy phosphate transfer in excitable tissues, thereby playing a central role in myocardial energy homeostasis. Creatine kinase knockout mice have been reported to exhibit left ventricular hypertrophy and dilation when assessed by cardiac magnetic resonance imaging (Nahrendorf et al., 2005). Consistent with the negative logFC observed in the D7 vs D0 comparison, IPA predicts that ventricular and myocardial functions may be compromised.

A similar pattern was observed for CSF1, a pro-inflammatory cytokine that regulates multiple aspects of the monocytic lineage, including the proliferation and differentiation of monocyte progenitors. In the context of post–myocardial infarction repair, CSF1 has been associated with reduced infarct size and scar formation, potentially through enhanced macrophage infiltration and neovascularization within the infarcted tissue (Morimoto et al., 2007). Despite its negative logFC in our dataset, IPA linked CSF1 to increased predicted activation of the cardiac abnormalities listed above.

CTSA encodes a lysosomal protease with broad substrate specificity. In studies of the secretome of adult mouse cardiac fibroblasts, superoxide dismutase was identified as a novel CTSA target following proteolytic processing. In transgenic mice overexpressing CTSA, impaired antioxidant activity was associated with increased superoxide radicals, inflammation, and myocyte hypertrophy, among other pathological processes (Hohl et al., 2020). In line with these observations, IPA associated the positive logFC for CTSA with an increased predicted state for cardiac complications.

Finally, for TNFRSF1B (TNFR2), IPA indicated that decreased expression between conditions was associated with an increased predicted state for cardiac disease. This directionality is supported by evidence from a Tnf^ΔARE^ mouse model, in which deletion of TNFRSF1B was linked to spondyloarthritis and heart valve stenosis, and where TNFRSF1B was implicated in the regulation of pathways related to proliferation, inflammation, migration, and other disease-relevant gene programs (Sakkou et al., 2018).

Collectively, these results highlight a rapid and multidimensional proteomic shift following TCZ administration, ranging from a hyperinflammatory and catabolic profile to one indicative of metabolic, endothelial, and immune recovery. The trajectories observed for key molecules such as SERPINA4, HDL-associated apolipoproteins, and IGKV3-11 not only reflect therapeutic response but may serve as pharmacodynamic markers and prognostic indicators for early cytokine-blockade interventions (Supplementary Figure 2) (Demichev et al., 2021; Filbin et al., 2021). Nevertheless, TCZ should be used with clinical caution in patients with high cardiovascular risk, especially those with baseline dyslipidemia, as IL-6 receptor blockade is consistently associated with early increases in total cholesterol, LDL cholesterol, and triglycerides, which warrants monitoring and control of baseline and treatment-related lipids according to guidelines (Pierini et al., 2021).

However, comparative safety data from rheumatoid arthritis do not show an excessive risk of serious adverse cardiovascular events with TCZ compared to tumor necrosis factor inhibitors or other biologic drugs; in fact, a large network meta-analysis and cohort studies with multiple databases report a similar cardiovascular risk, or in some analyses even numerically lower, than key comparators (including a possible reduction in myocardial infarction versus certain agents) (Castagné et al., 2019; Kim et al., 2017). In practice, our findings support careful cardiovascular and lipid monitoring rather than blanket exclusion of TCZ in individuals with high baseline risk.

## Conclusion

TCZ administration induced an early transition toward a proteomic phenotype associated with less severe disease in patients with COVID-19 pneumonia. These findings provide insight into the broad, multifaceted biological effects of TCZ and how they potentially modulate the clinical trajectory of severe COVID-19.

## Author Contributions

N.P.B. and J.R.: Study design, data collection, analysis and interpretation. J.C., G.B., P.H.S., B.L. and E.R.: data analysis and interpretation. P.S., M.H., C.V. and E.N-L.: data collection, analysis and interpretation. V.L., D.V. and E.S.K.: data interpretation. J.C., G.B., P.H.S., V.L. and N.P.B: writing of the drafted manuscript. All authors: revision of the drafted manuscript and approval of its final version.

## Acknowledgments

N.P.B. and J.R. were supported by a grant from the Pontificia Universidad Católica de Chile Interdisciplinary Seed Fund #I180031, J.R. was supported by ANID Fondecyt Regular grant #1241897, and N.P.B. was supported by ANID Fondecyt Regular grant #1251222 and ANID Fondequip grants #EQM150102 and #EQM170172.

