## Supplementary figures and images for "Tocilizumab induces significant changes in longitudinal proteomes of blood serum from patients with severe COVID-19 pneumonia"

### Suplementary Figure 2

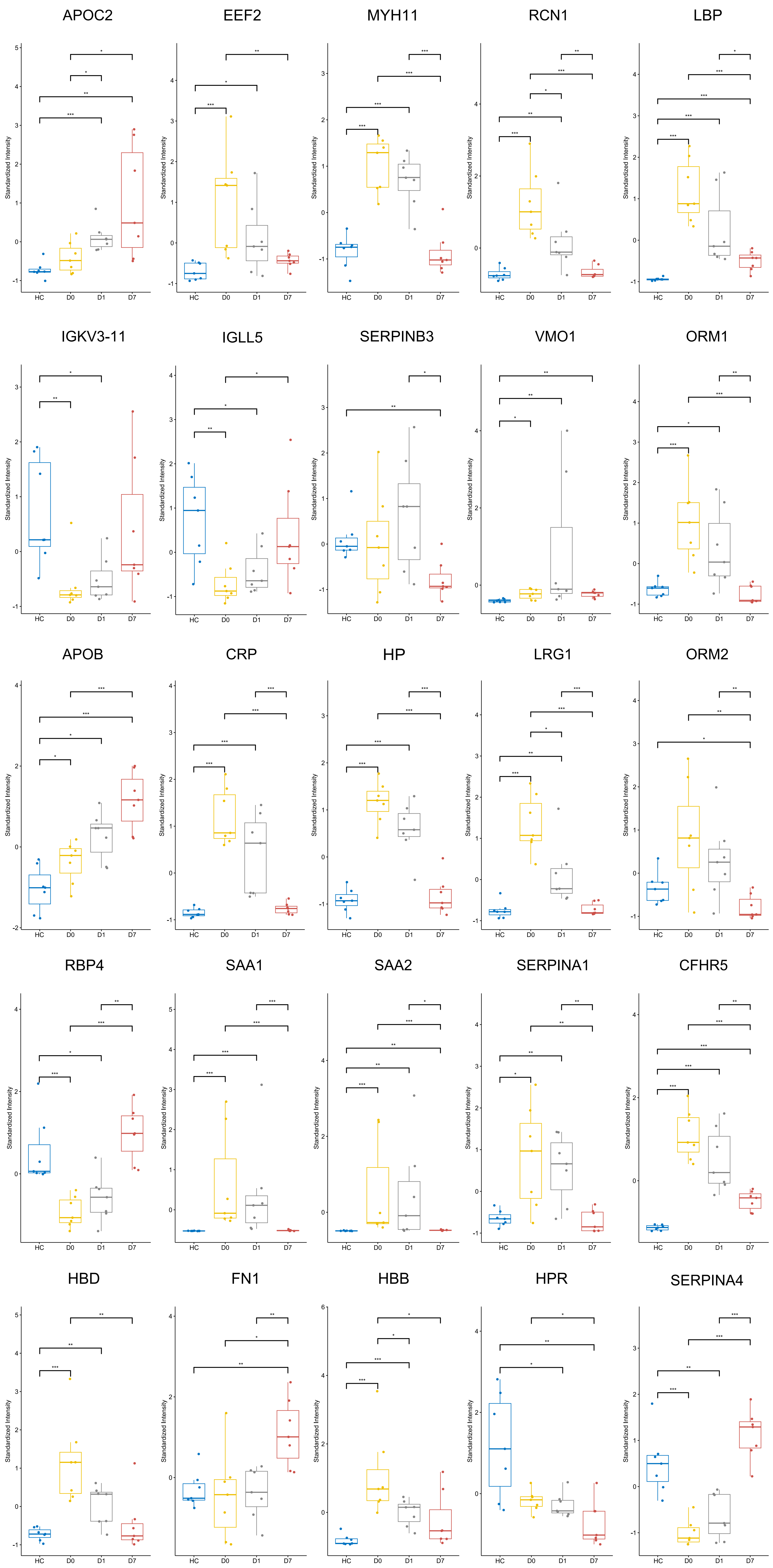

### Supplementary Figure 1

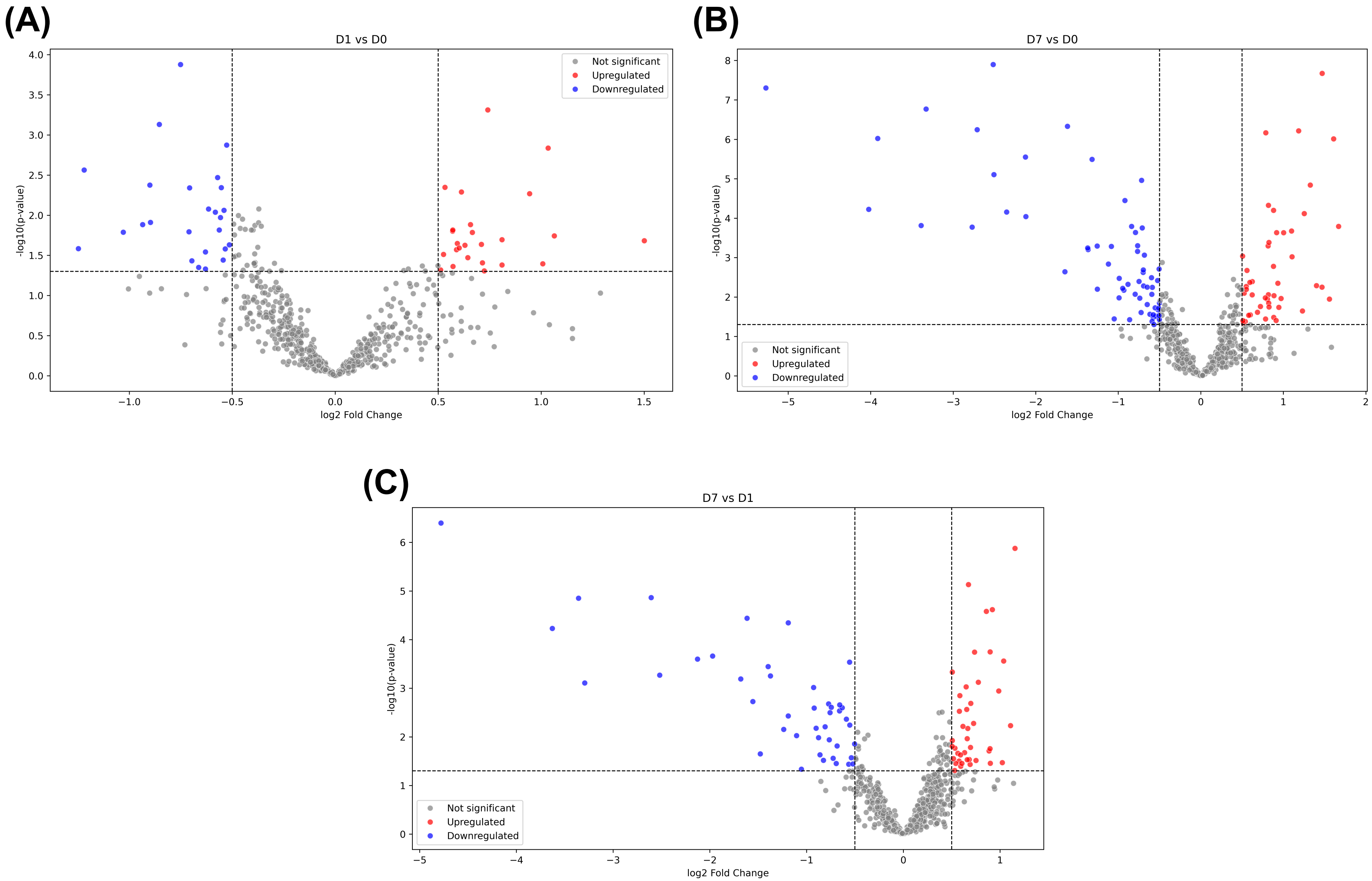

### Supplementary Figure 3

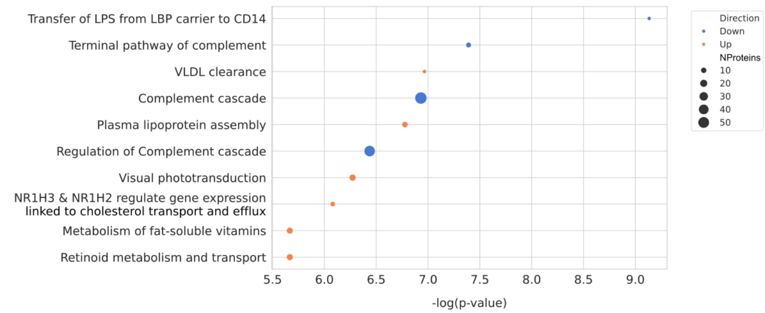
