## Supplementary Figure Legends for "Tocilizumab induces significant changes in longitudinal proteomes of blood serum from patients with severe COVID-19 pneumonia"

**Supplementary Figure 1. Volcano plots from severe COVID-19 pneumonia patients’ data.** (A) D1 vs D7, (B) D7 vs D0 and (C) D7 vs D1 volcano plot. Proteins were classified as up-regulated when the log₂ fold-change (log₂FC) > 0.5 and down-regulated when log₂FC < 0.5.

**Supplementary Figure 2. Box plots of top 25 DEPs in severe COVID-19 pneumonia patients.** Statistical comparisons were analyzed via Wilcoxon rank-sum test. n = 7; **p<0.05, **p<0.01, ***p<0.001.*

**Supplementary Figure 3. Reactome pathway enrichment analysis of D7 vs D1.** Bubble plot showing significantly enriched pathways identified from the protein expression dataset from the comparison between D7 vs D1. The x-axis represents statistical significance as −log(p-value), with higher values indicating stronger enrichment. The y-axis lists the enriched pathways. Bubble size corresponds to the number of proteins (NProteins) associated with each pathway, and color indicates directionality of regulation (orange: upregulated; blue: downregulated). Pathways related to complement activation, lipid metabolism, and vitamin/retinoid processing are prominently enriched.
